# Motor Cortical Gamma Organisation Following Preterm Birth Relates to Early Motor Development

**DOI:** 10.64898/2025.12.10.693534

**Authors:** Jaeyoung Yang, Shrisha Sathishkumar, Yusuke Mitani, Chiaki Hasegawa, Takashi Ikeda, Sanae Tanaka, Hong-gi Yeom, Kin Tung Jonathan Chan, Alice Waitt, Ken Yaoi, Sumie Iwasaki, Daisuke N. Saito, Tetsu Hirosawa, Mitsuru Kikuchi, Yuko Yoshimura, Kyung-min An

## Abstract

Preterm birth, defined as delivery before 37 weeks of gestation, is associated with differences in neurodevelopment and an increased likelihood of motor challenges that may persist throughout childhood. However, the neurophysiological correlates of these motor differences remain insufficiently understood. Oscillatory activity in the primary motor cortex is closely linked to motor execution, yet motor-related oscillations have rarely been examined in children born preterm. In this cross-sectional observational study, we investigated motor-related gamma oscillations in 5- to 7-year-old children using a child-customised magnetoencephalography system. Nineteen children born full-term and 18 children born preterm of comparable age performed a child-friendly dominant-hand finger movement task. Movement-related gamma activity was quantified from bilateral precentral gyrus regions used as anatomical proxies for the primary motor cortices. The preterm group showed longer mean response times and greater intra-individual response-time variability than the full-term group (*P* = 0.003 and *P* = 0.030, respectively). Contralateral gamma power was also lower in the preterm group (full-term: 59.89 ± 28.87%; preterm: 37.97 ± 25.82%; *t*(35) = 2.43, *P* = 0.020, Cohen’s *d* = 0.80). A significant Group × Hemisphere interaction (*F*(1, 35) = 13.36, *P* < 0.001, partial η² = 0.276) further indicated differences in the hemispheric organisation of gamma activity between groups. The gamma laterality index was also lower in children born preterm (full-term: 0.24 ± 0.21; preterm: −0.11 ± 0.37; *t*(26.4) = 3.51, *P* = 0.002, Cohen’s *d* = 1.17), consistent with reduced contralateral dominance. Across the whole sample, greater contralateral gamma lateralisation was associated with higher scores on the Hand Movements subtest of the Kaufman Assessment Battery for Children (ρ = 0.40, *P* = 0.014, false discovery rate-adjusted *q* = 0.042) and earlier independent walking (ρ = −0.57, *P* < 0.001, *q* = 0.002). The association with independent walking was also observed within the preterm group (ρ = −0.78, *P* < 0.001, *q* = 0.002). These findings suggest that reduced contralateral gamma power and reduced hemispheric lateralisation in children born preterm may reflect differences in the developmental organisation of motor cortical networks. This study provides new insight into motor cortical organisation and its relationship with motor development following preterm birth.

## Introduction

Preterm birth is defined as birth before 37 weeks of gestation.^1^ A global analysis estimated that 9.9% of infants born in 2020 were preterm.^2^ Preterm birth is associated with neurodevelopmental differences across multiple domains, including attention,^3–5^ cognition,^6–8^ motor function,^8–11^ and socio-emotional development.^12,13^ These neurodevelopmental differences can remain evident into school age and may affect later functional and academic outcomes.^14,15^ Evidence further suggests that the extent of these neurodevelopmental differences increases with lower gestational age.^16^

In terms of motor function, children born preterm show differences in gross and fine motor performance compared with children born full-term, and these differences can persist across development. Indeed, differences in motor performance have been observed as late as 8–10 years of age,^11^ and remain evident into adolescence.^17^ Furthermore, motor performance has been shown to be significantly associated with long-term cognitive function, academic achievement, and behavioural outcomes.^17,18^ Early motor development can also differ following preterm birth, including in the timing of foundational developmental milestones such as independent walking.^10^ Such milestones provide an important developmental context for investigating whether early motor history is related to the later functional organisation of the motor system. Together, these findings suggest that motor differences associated with preterm birth may reflect underlying differences in the maturation of neural systems supporting motor control. Identifying neurophysiological features associated with these developmental differences may therefore provide insight into motor-system development following preterm birth.

Neural oscillations represent rhythmic patterns of neural activity, and motor processes are known to modulate characteristic oscillatory rhythms. In particular, motor-related gamma oscillations, typically observed in the high-frequency range of approximately 70–90 Hz, emerge around movement onset and have been linked to motor initiation and execution.^19–22^ Studies in healthy adults have consistently demonstrated robust increases in gamma-band power in the contralateral motor cortex during unilateral movement, consistent with a predominantly contralateral organisation of movement-related gamma activity.^19–21^ Movement-related gamma activity has been observed in both children and adults,^19,21,23–25^ while the developmental organisation of these responses remains incompletely understood. Importantly, motor-related gamma power and its hemispheric distribution may provide complementary information about motor cortical function. Gamma power may reflect the magnitude of movement-related oscillatory activity, whereas the degree of hemispheric lateralisation may index the relative hemispheric organisation of the motor network. Examining these measures together may therefore distinguish differences in the magnitude of motor-related cortical activity from differences in its functional organisation. Differences in movement-related gamma activity have also been reported in individuals with autism spectrum disorder,^23,24,26^ suggesting that motor-related gamma measures may be sensitive to variation in motor cortical organisation. However, whether motor-related gamma activity differs following preterm birth remains unclear. In particular, little is known about whether the magnitude and hemispheric lateralisation of motor-related gamma activity differ between children born full-term and preterm, or whether individual variation in gamma lateralisation relates to motor development.

Preterm birth may affect the development of cortical and white matter networks that support coordinated motor processing, with previous neuroimaging studies reporting differences in white matter development, thalamocortical pathways, and interhemispheric structures following preterm birth.^27–32^ Recent functional MRI evidence in 6-year-old children born very preterm has also demonstrated persistent differences in task-evoked motor activation and large-scale resting-state functional organisation.^33^ Examining movement-related gamma power and its hemispheric lateralisation may therefore provide a functional approach to characterising motor cortical organisation following preterm birth. Assessing both the magnitude and hemispheric distribution of gamma activity may further clarify whether individual differences in motor-network organisation are related to developmental motor history.

In the present study, we hypothesised that children born preterm would show differences in movement-related gamma oscillations, in terms of both power and hemispheric lateralisation, when compared with children born full-term. Specifically, we predicted reduced contralateral gamma power and reduced hemispheric lateralisation in children born preterm. We further predicted that the preterm group would demonstrate longer mean response times and greater intra-individual response-time variability during the motor task, together with group differences in standardised motor measures. Finally, we hypothesised that greater contralateral gamma lateralisation would be associated with better motor performance and earlier attainment of motor developmental milestones.

To test these hypotheses, we measured motor-related cortical oscillations during a child-friendly, video game–like unilateral finger movement task. Data were acquired using a child-customised magnetoencephalography (MEG) system, which provides high temporal resolution, and source-level analyses were used to characterise cortical activity. This approach allowed us to characterise motor-related gamma activity and its hemispheric organisation in 5- to 7-year-old children born full-term and preterm.

## Materials and Methods

### Participants

The final sample comprised 37 children, including 19 children born full-term (mean age ± SD, 69.05 ± 6.92 months; six females) and 18 children born preterm (69.28 ± 3.64 months; six females). All participants were right-handed according to the Edinburgh Handedness Inventory.^34^ Participants were recruited and assessed between 2017 and 2019 through community- and hospital-based routes in Kanazawa and the surrounding area, Japan. No formal a priori sample-size calculation was performed. The recruitment target was informed by the sample sizes of previous paediatric MEG studies using the same or closely comparable motor paradigm, which included approximately 14–19 children per group.^23,24^ A total recruitment target of 40 children was therefore set to obtain group sizes comparable with these previous studies while allowing for anticipated participant loss due to non-attendance or unsuccessful MEG/MRI acquisition. Three children (one born full-term and two born preterm) were not included in the final sample because MEG and/or MRI acquisition could not be initiated on the day of assessment.

Full-term participants were recruited from the local community through advertisements distributed via nursery schools, primary schools, the university community, the paediatric network affiliated with Kanazawa University Hospital, and local public health centres. Birth and medical histories were screened to determine eligibility. Full-term participants were born between 38 and 42 weeks of gestation, had no history of perinatal complications, and had no known neurological or developmental disorders.

Children born preterm were recruited through collaborating paediatricians at Kanazawa University Hospital, who reviewed their neonatal and medical records. Children born before 37 weeks of gestation were included in the preterm group. To minimise potential confounding effects of major neurological or structural brain abnormalities, exclusion criteria were applied based on clinical history. Children with a documented medical history of severe white matter injury, periventricular leukomalacia, grade III/IV intraventricular haemorrhage, cerebral palsy, or any known gross structural brain abnormalities were excluded from the study.

Parents of all children provided written informed consent to participate in the study, and all procedures were approved by the Ethics Committee of Kanazawa University Hospital in accordance with the Declaration of Helsinki. Gestational age at birth was obtained from parental reports and verified against available birth or medical records. Age at experiment did not differ between the full-term and preterm groups (full-term: 69.05 ± 6.92 months; preterm: 69.28 ± 3.64 months; *t*(27.6) = −0.13, *P* = 0.902, Cohen’s *d* = −0.04), and Achievement scores on the Kaufman Assessment Battery for Children (K-ABC)^35^ also did not differ significantly between groups (full-term: 106.00 ± 15.79; preterm: 98.17 ± 16.91; *t*(35) = 1.46, *P* = 0.154, Cohen’s *d* = 0.48; **Table 1**). Gestational age, birth weight, birth length and birth head circumference were significantly lower in the preterm group (all *P* < 0.001).

**Table 1.**
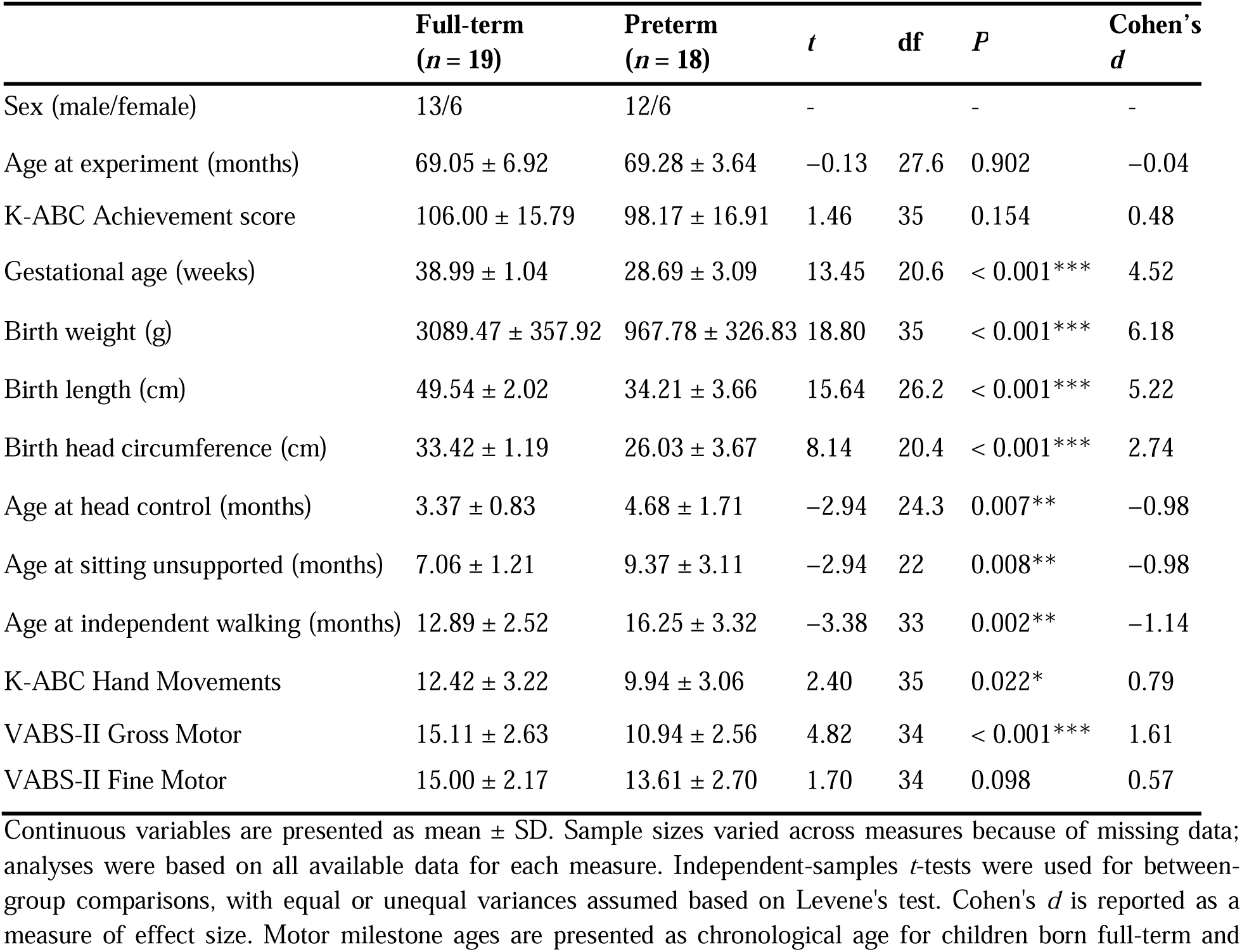
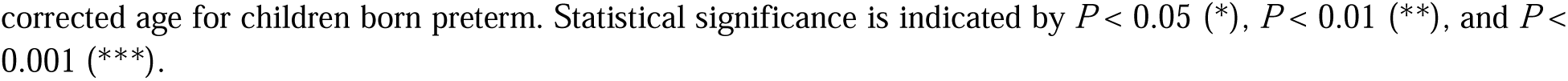
Participant characteristics.

|  | Full-term<br>(*n* = 19) | Preterm<br>(*n* = 18) | *t* | df | *P* | Cohen's<br>*d* |
| --- | --- | --- | --- | --- | --- | --- |
| Sex (male/female) | 13/6 | 12/6 | - | - | - | - |
| Age at experiment (months) | $69.05 \pm 6.92$ | $69.28 \pm 3.64$ | -0.13 | 27.6 | 0.902 | -0.04 |
| K-ABC Achievement score | $106.00 \pm 15.79$ | $98.17 \pm 16.91$ | 1.46 | 35 | 0.154 | 0.48 |
| Gestational age (weeks) | $38.99 \pm 1.04$ | $28.69 \pm 3.09$ | 13.45 | 20.6 | < 0.001*** | 4.52 |
| Birth weight (g) | $3089.47 \pm 357.92$ | $967.78 \pm 326.83$ | 18.80 | 35 | < 0.001*** | 6.18 |
| Birth length (cm) | $49.54 \pm 2.02$ | $34.21 \pm 3.66$ | 15.64 | 26.2 | < 0.001*** | 5.22 |
| Birth head circumference (cm) | $33.42 \pm 1.19$ | $26.03 \pm 3.67$ | 8.14 | 20.4 | < 0.001*** | 2.74 |
| Age at head control (months) | $3.37 \pm 0.83$ | $4.68 \pm 1.71$ | -2.94 | 24.3 | 0.007** | -0.98 |
| Age at sitting unsupported (months) | $7.06 \pm 1.21$ | $9.37 \pm 3.11$ | -2.94 | 22 | 0.008** | -0.98 |
| Age at independent walking (months) | $12.89 \pm 2.52$ | $16.25 \pm 3.32$ | -3.38 | 33 | 0.002** | -1.14 |
| K-ABC Hand Movements | $12.42 \pm 3.22$ | $9.94 \pm 3.06$ | 2.40 | 35 | 0.022* | 0.79 |
| VABS-II Gross Motor | $15.11 \pm 2.63$ | $10.94 \pm 2.56$ | 4.82 | 34 | < 0.001*** | 1.61 |
| VABS-II Fine Motor | $15.00 \pm 2.17$ | $13.61 \pm 2.70$ | 1.70 | 34 | 0.098 | 0.57 |

Continuous variables are presented as mean ± SD. Sample sizes varied across measures because of missing data; analyses were based on all available data for each measure. Independent-samples *t*-tests were used for between-group comparisons, with equal or unequal variances assumed based on Levene's test. Cohen's *d* is reported as a measure of effect size. Motor milestone ages are presented as chronological age for children born full-term and
corrected age for children born preterm. Statistical significance is indicated by $P < 0.05$ (\*), $P < 0.01$ (\*\*), and $P < 0.001$ (\*\*\*)

Ages at attainment of early motor milestones, including head control, sitting unsupported, and independent walking, were obtained from records in the Japanese Maternal and Child Health Handbook (Boshi Kenko Techo). For children born full-term, milestone ages were expressed as chronological age from birth, whereas for children born preterm, milestone ages were expressed as corrected age, accounting for gestational age at birth. Corrected age was calculated relative to 40 weeks of gestation. Children born preterm attained head control (full-term: 3.37 ± 0.83 months; preterm: 4.68 ± 1.71 months; *t*(24.3) = −2.94, *P* = 0.007, Cohen’s *d* = −0.98), sitting unsupported (full-term: 7.06 ± 1.21 months; preterm: 9.37 ± 3.11 months; *t*(22) = −2.94, *P* = 0.008, Cohen’s *d* = −0.98), and independent walking (full-term: 12.89 ± 2.52 months; preterm: 16.25 ± 3.32 months; *t*(33) = −3.38, *P* = 0.002, Cohen’s *d* = −1.14) at significantly older ages than children born full-term (**Table 1**).

Motor-related performance was evaluated using the Hand Movements subtest of the K-ABC and the Gross Motor and Fine Motor subdomains of the Motor Skills Domain of the Vineland Adaptive Behaviour Scales, Second Edition (VABS-II).^36^ The preterm group showed significantly lower K-ABC Hand Movements scores than the full-term group (full-term: 12.42 ± 3.22; preterm: 9.94 ± 3.06; *t*(35) = 2.40, *P* = 0.022, Cohen’s *d* = 0.79; **Table 1**), whereas the difference in VABS-II Fine Motor scores did not reach statistical significance (full-term: 15.00 ± 2.17; preterm: 13.61 ± 2.70; *t*(34) = 1.70, *P* = 0.098, Cohen’s *d* = 0.57). In contrast, VABS-II Gross Motor scores were significantly lower in the preterm group (full-term: 15.11 ± 2.63; preterm: 10.94 ± 2.56; *t*(34) = 4.82, *P* < 0.001, Cohen’s *d* = 1.61). The K-ABC Hand Movements and VABS-II Gross Motor scores are also shown in **Fig. 1B**. Additional demographic and behavioural details are provided in **Table 1**.

**Figure 1.**
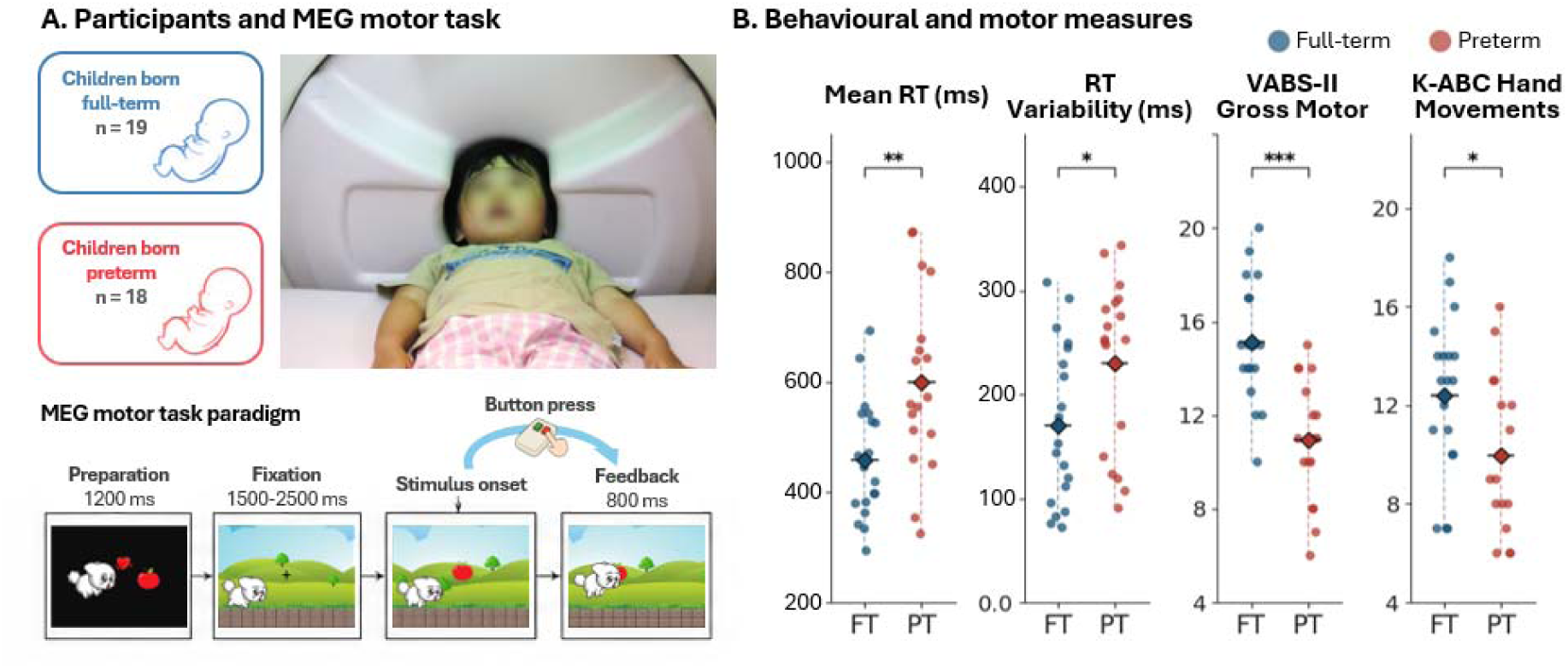
MEG experimental paradigm and behavioural measures in children born full-term and preterm. **(A) Participants and MEG motor task.** The task involved a child-friendly video game in which a cartoon puppy jumped to catch a target fruit that appeared at the fixation point. Participants pressed a button upon visual target onset, causing the puppy to jump and retrieve the fruit. Following a correct button press, an 800-ms feedback animation showed the puppy catching the target fruit. A separate reward animation was presented after each block to maintain motivation. The task paradigm was based on An et al.^23,24^ **(B) Behavioural and motor measures.** Childre born preterm showed longer response times and greater intra-individual response-time variability than children bor full-term. Mean response time was significantly longer in the preterm group (full-term: 459 ± 108 ms; preterm: 601 ± 162 ms; *t*(35) = −3.14, *P* = 0.003, Cohen’s *d* = −1.03). Intra-individual response-time variability was also significantly greater in the preterm group (full-term: 171 ± 77 ms; preterm: 230 ± 82 ms; *t*(35) = −2.27, *P* = 0.030, Cohen’s *d* = −0.75). VABS-II Gross Motor scores were significantly lower in the preterm group (full-term: 15.11 ± 2.63; preterm: 10.94 ± 2.56; *t*(34) = 4.82, *P* < 0.001, Cohen’s *d* = 1.61). K-ABC Hand Movements scores were also significantly lower in the preterm group (full-term: 12.42 ± 3.22; preterm: 9.94 ± 3.06; *t*(35) = 2.40, *P* = 0.022, Cohen’s *d* = 0.79). Dots represent individual participants and diamonds indicate group means. Two-tailed independent-samples *t*-tests were used for the comparisons shown in panel B. Mean response time, response-time variability, and K-ABC Hand Movements included 19 children born full-term and 18 children born preterm; VABS-II Gross Motor included 18 children in each group. Each data point represents one child. RT, response time; FT, full-term; PT, preterm. Statistical significance is indicated by *P* < 0.05 (*), *P* < 0.01 (**), and *P* < 0.001 (***). **Alt text:** Two-panel figure illustrating the paediatric MEG motor task and behavioural results. Panel A shows children born full-term and preterm, a child positioned in the paediatric MEG system, and the sequence of the video game task, in which a preparation cue and fixation period are followed by visual target onset, a right index-finger button press, and feedback. Panel B shows individual participant data for mean response time, response-time variability, VABS-II Gross Motor scores, and K-ABC Hand Movements scores. Children born preterm show longer and more variable response times and lower Gross Motor and Hand Movements scores than children born full-term.

### Experimental design

We employed a unilateral finger movement video game paradigm (**Fig. 1A**) developed in our previous studies.^23,24^ The task was programmed using Presentation software (Neurobehavioral Systems) and designed to resemble a child-friendly game. Participants lay supine throughout the session to facilitate comfort and minimise head movement.

At the beginning of each trial, a “mission” cue appeared on the screen, indicating the target fruit (e.g. an apple or a grape) that the cartoon puppy would attempt to collect. After 1200 ms, the puppy appeared on the left side of the display while a central fixation cross was presented to maintain gaze and reduce eye movement artefacts. After a random delay of 1.5–2.5 s, the fixation cross was replaced by the target fruit image, which served as the go signal. Participants were instructed to press a button using their dominant (right) index finger as quickly as possible upon target appearance.

When a correct response was made, the puppy jumped and caught the fruit for 800 ms; this was followed by a brief inter-trial interval of 3.5–4.5 s before the next trial began. If a participant pressed the button prematurely or failed to respond, the puppy fell down and the trial was repeated. Each block comprised 10 successfully completed trials, and participants completed 10 blocks, yielding 100 successfully completed trials in total. After each block, a short “reward” animation was presented in which the puppy received a bone and a fanfare sound, helping maintain attention and motivation. Prior to data collection, participants completed one practice block to familiarise themselves with the task.

### Magnetoencephalography recording

MEG signals were recorded with a child-customised 151-channel MEG system (PQ51R; Yokogawa/KIT, Kanazawa, Japan) in a magnetically shielded room. To support participants’ comfort during the MEG measurements, two staff members remained in the shielded room throughout the recording, and parents were able to monitor their child’s progress via a television screen. No recording was terminated because of participant distress or discomfort. Stimuli were projected onto a rear-projection screen using an LCD projector (IPSiO PJWX6170N, Ricoh), with a visual angle of approximately 21° (vertical) × 26° (horizontal). Responses were collected via a nonmagnetic fibre-optic button box (LUMINA LSC400 with LS-PAIR response box, Cedrus Corporation). Before recording, four head-position coils were attached at Cz, 5 cm anterior to Cz, and 5 cm superior to each of the left and right preauricular points to determine head position within the MEG helmet. The locations of the coils and 100 scalp surface points were digitised using a 3D digitiser (FastSCAN, Polhemus).

During recording, MEG signals were sampled at 2000 Hz with a 200 Hz low-pass filter. Following MEG acquisition, the coils were replaced with MRI-visible markers, and T1-weighted structural MRI images were obtained for each participant using a 1.5 T SIGNA Explorer system (GE Healthcare) with a T1-weighted gradient-echo Silenz pulse sequence. MRI parameters were as follows: repetition time (TR) = 435.68 ms, echo time (TE) = 0.024 ms, flip angle = 7°, field of view (FOV) = 220 mm, matrix size = 256 × 256, slice thickness = 1.7 mm, number of slices = 130. The anatomical images were used for MEG–MRI co-registration during source localisation analysis.

### Data analysis

MEG data preprocessing was performed using Brainstorm.^37^ The data were down-sampled to 500 Hz and band-pass filtered between 0.5 and 100 Hz. A notch filter with a 1-Hz bandwidth was applied at 60 Hz to attenuate power-line noise. Bad channels, including flat and excessively noisy channels, were identified by visual inspection, marked as bad, and excluded from subsequent analyses.

To remove non-neural artefacts, including eye blinks, eye movements, and cardiac activity, independent component analysis (ICA) was performed on the continuous preprocessed MEG data using the Infomax algorithm,^38,39^ with 20 independent components. Components associated with ocular and cardiac artefacts were identified by visual inspection of their sensor topographies and component time courses and were removed. The remaining components were then back-projected to reconstruct the cleaned MEG signals.

The preprocessed continuous data were epoched from −3 to 3 s relative to button-press onset. Only trials with a valid button press occurring 200–2000 ms after visual target onset were retained; trials with no response or premature or accidental responses were excluded. Epochs containing visually identified muscle artefacts within the −2 to 2 s window relative to button-press onset were excluded from further analysis.

### Source estimation

Source estimation was performed on the preprocessed epochs using weighted minimum-norm estimation^40^ implemented in the Brainstorm toolbox. MEG data were co-registered with individual MRI scans to enable source localisation. An overlapping-spheres head model was constructed for each participant using their individual MRI anatomy. The noise covariance matrix was estimated from the baseline period (−2000 to −1500 ms relative to button-press onset). Tikhonov regularisation was applied with a regularisation parameter of λ = 0.1.

To investigate motor-related activity in the precentral gyrus, source time courses were extracted from the bilateral precentral gyri, corresponding to the left and right “Precentral” anatomical parcellations of the Desikan-Killiany atlas.^41^ These precentral gyrus regions were used as anatomical proxies for the primary motor cortices (M1). For the right index-finger movement, the left precentral gyrus was defined as contralateral M1 and the right precentral gyrus as ipsilateral M1. To obtain a representative M1 time course from the spatially distributed source activity, principal component analysis (PCA) was applied to the source time courses within each precentral gyrus. The first principal component (PC1), which captured the greatest variance in the regional source activity, was used for subsequent time-frequency analyses (**Fig. 2A**).

**Figure 2.**
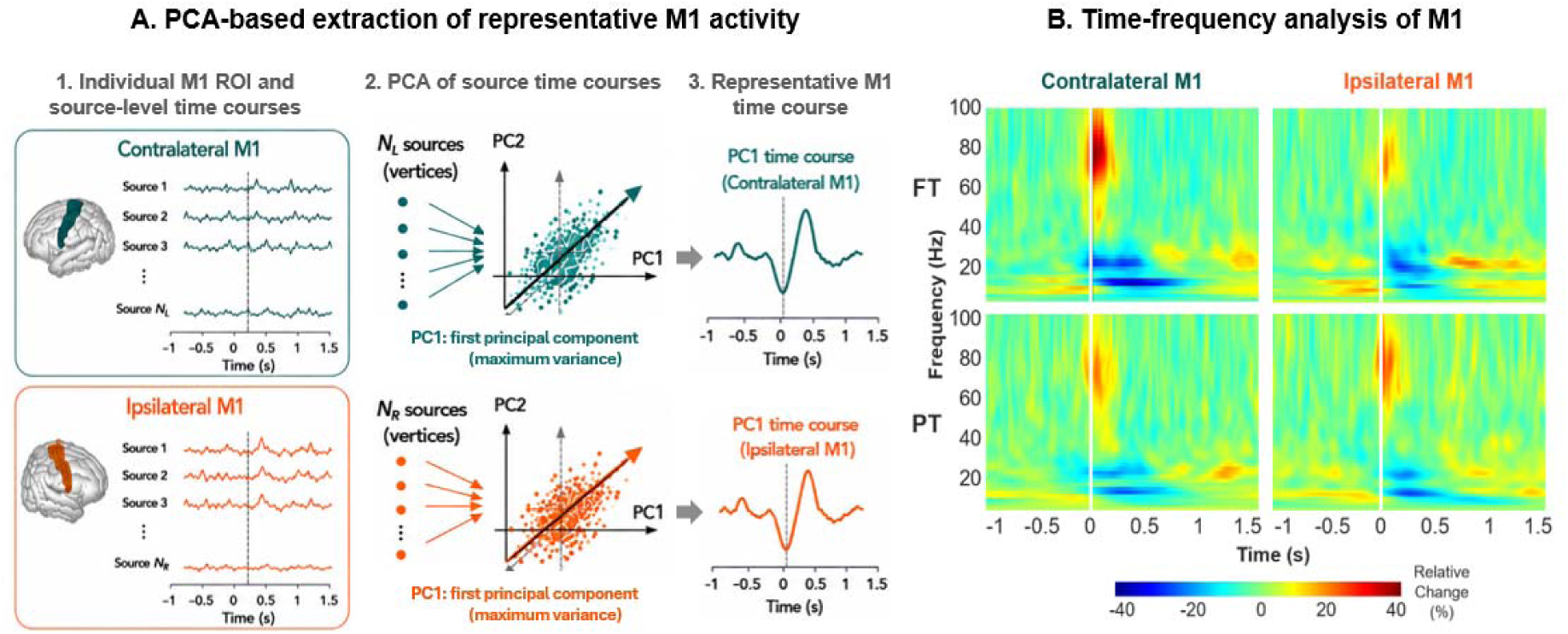
Time-frequency representations of M1 activity in children born full-term and children born preterm. Source-level time courses were extracted from the left and right precentral gyrus anatomical parcellations, which were used as anatomical proxies for M1. Principal component analysis (PCA) was then applied to derive a representative time course capturing the dominant pattern of variance within each region. **(A)** Schematic illustration of the PCA-based extraction of representative M1 activity from contralateral and ipsilateral M1. **(B)** Grand-averaged time-frequency representations of the first principal component for contralateral and ipsilateral M1 in children born full-term and children born preterm. 0 ms denotes button-press onset. The plots illustrate the distribution of oscillatory activity across time and frequency for each group and hemisphere. Gamma-band activity was quantified at the participant level and compared between groups (see Fig. 3 and **Table 3**). Grand-averaged TFRs were calculated from 19 children born full-term and 18 children born preterm. M1, primary motor cortex; FT, full-term; PT, preterm. **Alt text:** Two-panel figure illustrating the extraction and time-frequency analysis of bilateral M1 activity. Panel A schematically shows source time courses from the contralateral and ipsilateral precentral gyrus regions, principal component analysis of the regional source activity, and extraction of the first principal component as the representative M1 time course. Panel B shows time-frequency maps for contralateral and ipsilateral M1 in children born full-term and preterm, with movement-related gamma activity in the 70–90 Hz range around button-press onset.

### Time-frequency analysis

The representative bilateral M1 source time courses were used to calculate time-frequency representations (TFRs) across the 1–100 Hz frequency range using a seven-cycle Morlet wavelet transformation.^42^ TFRs were baseline-normalised to express spectral power as a percentage change relative to the mean power during the baseline interval (−2000 to −1500 ms relative to button-press onset). TFRs were calculated separately for each participant and hemisphere, and participant-level TFRs were subsequently grand-averaged within each group for visualisation.

Based on previous work using the same motor paradigm,^23,24^ movement-related gamma power was quantified in the 70–90 Hz range and 0–100 ms following button-press onset. Gamma power was therefore quantified for each participant and hemisphere by averaging the baseline-normalised spectral power across the 70–90 Hz frequency range and 0–100 ms post-button-press window. Peak gamma frequency was additionally determined within the 70–90 Hz frequency range and 0–100 ms time window for each participant and hemisphere.

### Laterality index

To quantify the relative hemispheric dominance of movement-related gamma activity, a laterality index (LI) was calculated for each participant according to the following equation^43^:

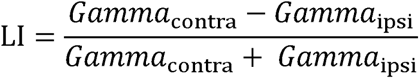

where Gamma and Gamma represent the mean baseline-normalised gamma power within the 70–90 Hz frequency range and 0–100 ms post-button-press window in the contralateral and ipsilateral M1, respectively. Positive LI values indicate greater contralateral than ipsilateral gamma power, whereas negative values indicate greater ipsilateral than contralateral gamma power. An LI close to zero indicates relatively balanced gamma power between the two hemispheres.

### Statistical analysis

Statistical analyses were conducted using Python (version 3.10) with the pandas, NumPy, SciPy, and Pingouin packages. Descriptive statistics, including means and standard deviations, were calculated for demographic, perinatal, behavioural, and neurophysiological variables. Group differences between children born full-term and children born preterm were assessed using two-tailed independent-samples *t*-tests. Levene’s test was used to assess the equality of variances, and Welch’s *t*-test was applied when the assumption of equal variances was violated. Cohen’s *d* was calculated as a measure of effect size.

To test whether the hemispheric distribution of movement-related gamma power differed between groups, a two-way mixed analysis of variance (ANOVA) was performed with group (full-term, preterm) as the between-subjects factor and hemisphere (contralateral, ipsilateral) as the within-subjects factor. A significant Group × Hemisphere interaction was followed by two-tailed paired-samples t-tests comparing contralateral and ipsilateral gamma power separately within the full-term and preterm groups.

Associations between behavioural response-time measures and perinatal characteristics and concurrent motor performance were examined using two-tailed Spearman’s rank correlation analyses, given the modest sample size and the potential influence of non-normal or skewed distributions. Mean response time and intra-individual response-time variability were each correlated with gestational age, birth weight, birth length, birth head circumference, K-ABC Hand Movements scores, VABS-II Gross Motor scores, and VABS-II Fine Motor scores. To account for multiple comparisons, Benjamini–Hochberg false discovery rate (FDR) correction was applied separately across the seven correlations examined for mean response time and the seven correlations examined for intra-individual response-time variability.

Associations between the gamma laterality index and early motor developmental milestones and concurrent motor performance were examined using two-tailed Spearman’s rank correlation analyses. Analyses were conducted across the whole sample and separately within the preterm group. Developmental and motor measures included age at head control, age at sitting unsupported, age at independent walking, K-ABC Hand Movements scores, VABS-II Gross Motor scores, and VABS-II Fine Motor scores. These analyses were conducted to test the hypothesis that greater contralateral gamma lateralisation would be associated with better motor performance and earlier attainment of motor developmental milestones. Benjamini–Hochberg FDR correction was applied separately across the six correlations examined in the whole sample and the corresponding six correlations examined within the preterm group.

For each correlation, Spearman’s correlation coefficient (ρ), uncorrected two-tailed *P*-value, FDR-adjusted *q*-value, and the number of participants included in the analysis (*n*) were reported. All statistical tests were two-tailed. For correlation analyses, statistical significance was defined as *q* < 0.05 following FDR correction; for all other statistical tests, statistical significance was set at *P* < 0.05.

## Results

### Behavioural response time and motor performance

Trials in which participants pressed the button within 200–2000 ms following visual target onset were included; trials with no response or premature or accidental responses were excluded. Button response time was defined as the latency between visual target onset and button-press onset (**Fig. 1A**). For each participant, mean response time was calculated as the mean of response times across all valid trials. The full-term group had a shorter mean response time than the preterm group (mean ± SD: full-term, 459 ± 108 ms; preterm, 601 ± 162 ms; *t*(35) = −3.14, *P* = 0.003; **Fig. 1B**). To assess trial-to-trial response consistency, we calculated the intra-individual standard deviation of response times for each participant. Intra-individual response-time variability was significantly lower in the full-term group than in the preterm group (mean ± SD: full-term, 171 ± 77 ms; preterm, 230 ± 82 ms; *t*(35) = −2.27, *P* = 0.030; **Fig. 1B**), indicating greater trial-to-trial variability in the preterm group.

Additionally, we examined associations between response-time measures and perinatal and motor performance measures (**Table 2**). After FDR correction, mean response time was negatively associated with gestational age (*ρ* = −0.44, *P* = 0.006, *q* = 0.008, *n* = 37), birth weight (ρ = −0.48, *P* = 0.004, *q* = 0.007, *n* = 37), birth length (ρ = −0.49, *P* = 0.002, *q* = 0.005, *n* = 37), birth head circumference (ρ = −0.37, *P* = 0.024, *q* = 0.024, *n* = 37), K-ABC Hand Movements scores (*ρ* = −0.55, *P* < 0.001, *q* = 0.003, *n* = 37), VABS-II Gross Motor scores (ρ = −0.40, *P* = 0.016, *q* = 0.019, *n* = 36), and VABS-II Fine Motor scores (ρ = −0.51, *P* = 0.002, *q* = 0.005, *n* = 36). Thus, longer response times were associated with lower gestational age, birth weight, birth length, and birth head circumference, as well as lower motor performance scores.

**Table 2.**
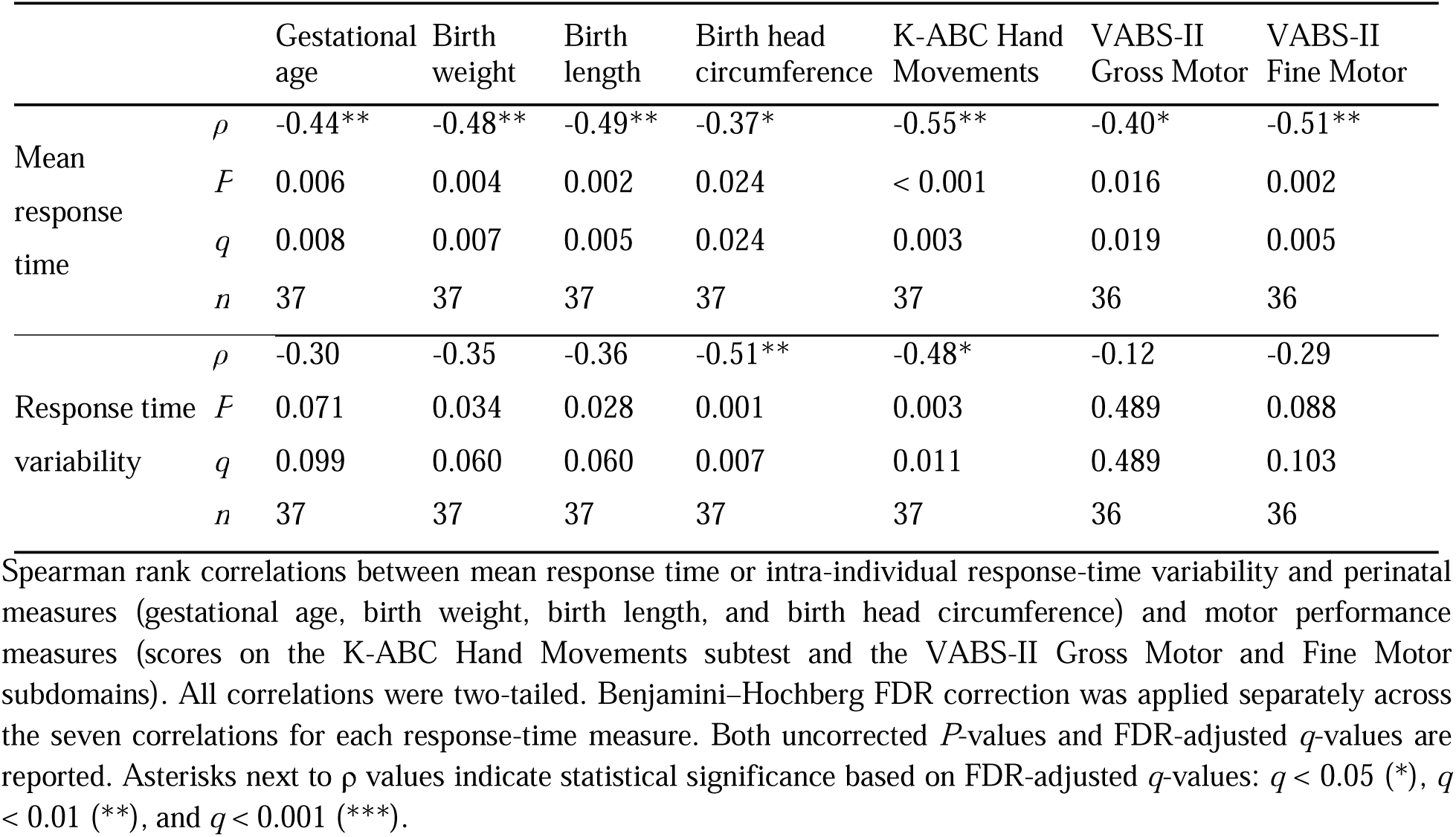
Correlations of response time measures with perinatal measures and motor performance scores.

|  |  | Gestational age | Birth weight | Birth length | Birth head circumference | K-ABC Hand Movements | VABS-II Gross Motor | VABS-II Fine Motor |
| --- | --- | --- | --- | --- | --- | --- | --- | --- |
| Mean response time | $\rho$ | -0.44** | -0.48** | -0.49** | -0.37* | -0.55** | -0.40* | -0.51** |
| | $P$ | 0.006 | 0.004 | 0.002 | 0.024 | < 0.001 | 0.016 | 0.002 |
| | $q$ | 0.008 | 0.007 | 0.005 | 0.024 | 0.003 | 0.019 | 0.005 |
| | $n$ | 37 | 37 | 37 | 37 | 37 | 36 | 36 |
| Response time variability | $\rho$ | -0.30 | -0.35 | -0.36 | -0.51** | -0.48* | -0.12 | -0.29 |
| | $P$ | 0.071 | 0.034 | 0.028 | 0.001 | 0.003 | 0.489 | 0.088 |
| | $q$ | 0.099 | 0.060 | 0.060 | 0.007 | 0.011 | 0.489 | 0.103 |
| | $n$ | 37 | 37 | 37 | 37 | 37 | 36 | 36 |

Spearman rank correlations between mean response time or intra-individual response-time variability and perinatal measures (gestational age, birth weight, birth length, and birth head circumference) and motor performance measures (scores on the K-ABC Hand Movements subtest and the VABS-II Gross Motor and Fine Motor subdomains). All correlations were two-tailed. Benjamini–Hochberg FDR correction was applied separately across the seven correlations for each response-time measure. Both uncorrected $P$ -values and FDR-adjusted $q$ -values are reported. Asterisks next to $\rho$ values indicate statistical significance based on FDR-adjusted $q$ -values: $q < 0.05$ (\*), $q < 0.01$ (\*\*), and $q < 0.001$ (\*\*\*).

**Table 3.** Comparison of M1 gamma activity between full-term and preterm groups.

|  | Full-term (*n* = 19) | Preterm (*n* = 18) | *t* | df | *P* | Cohen's *d* |
| --- | --- | --- | --- | --- | --- | --- |
|  | Mean (SD) | Mean (SD) |  |  |  |  |
| **Contralateral gamma oscillations** |  |  |  |  |  |  |
| Power (%) | 59.89 (28.87) | 37.97 (25.82) | 2.43 | 35 | 0.020* | 0.80 |
| Peak frequency (Hz) | 78.90 (5.56) | 76.67 (6.26) | 1.15 | 35 | 0.259 | 0.38 |
| **Ipsilateral gamma oscillations** |  |  |  |  |  |  |
| Power (%) | 36.03 (16.76) | 54.01 (46.62) | −1.55 | 21.1 | 0.137 | −0.52 |
| Peak frequency (Hz) | 77.32 (5.86) | 78.33 (6.40) | −0.51 | 35 | 0.617 | −0.17 |
| **Laterality index** | 0.24 (0.21) | −0.11 (0.37) | 3.51 | 26.4 | 0.002** | 1.17 |

Mean (SD) values for gamma power (%) and peak frequency (Hz) in the contralateral and ipsilateral M1 regions are shown for the full-term and preterm groups. The laterality index is also shown for each group. Independent-samples *t*-tests were used for between-group comparisons, with equal or unequal variances assumed based on Levene's test. Cohen's *d* is reported as a measure of effect size. Statistical significance is indicated by $P < 0.05$ (\*) and $P < 0.01$ (\*\*).

For intra-individual response-time variability, negative associations with birth head circumference (ρ = −0.51, *P* = 0.001, *q* = 0.007, *n* = 37) and K-ABC Hand Movements scores (*ρ* = −0.48, *P* = 0.003, *q* = 0.011, *n* = 37) remained significant after FDR correction. Associations with birth weight (*ρ* = −0.35, *P* = 0.034, *q* = 0.060, *n* = 37) and birth length (*ρ* = −0.36, *P* = 0.028, *q* = 0.060, *n* = 37) did not remain significant after FDR correction. No significant associations were observed with gestational age (ρ = −0.30, *P* = 0.071, *q* = 0.099, *n* = 37), VABS-II Gross Motor scores (*ρ* = −0.12, *P* = 0.489, *q* = 0.489, *n* = 36), or VABS-II Fine Motor scores (ρ = −0.29, *P* = 0.088, *q* = 0.103, *n* = 36).

### Motor-related gamma oscillations

Grand-averaged time–frequency representations (TFRs) from the bilateral precentral gyrus regions used as anatomical proxies for M1 during the finger-movement task are presented separately for the full-term and preterm groups (**Fig. 2B**). A motor-related gamma response was observed in both contralateral and ipsilateral M1 within the 70–90 Hz range around button-press onset.

Motor-related gamma power in the contralateral M1 was significantly higher in children born full-term (59.89 ± 28.87%) than in children born preterm (37.97 ± 25.82%), *t*(35) = 2.43, *P* = 0.020, Cohen’s *d* = 0.80 (**Fig. 3A**; **Table 3**). In contrast, ipsilateral M1 gamma power did not differ significantly between children born full-term (36.03 ± 16.76%) and children born preterm (54.01 ± 46.62%), *t*(21.1) = −1.55, *P* = 0.137, Cohen’s *d* = −0.52. No significant group differences were observed in peak gamma frequency in either the contralateral (*t*(35) = 1.15, *P* = 0.259) or ipsilateral M1 (*t*(35) = −0.51, *P* = 0.617) (**Table 3**).

**Figure 3.**
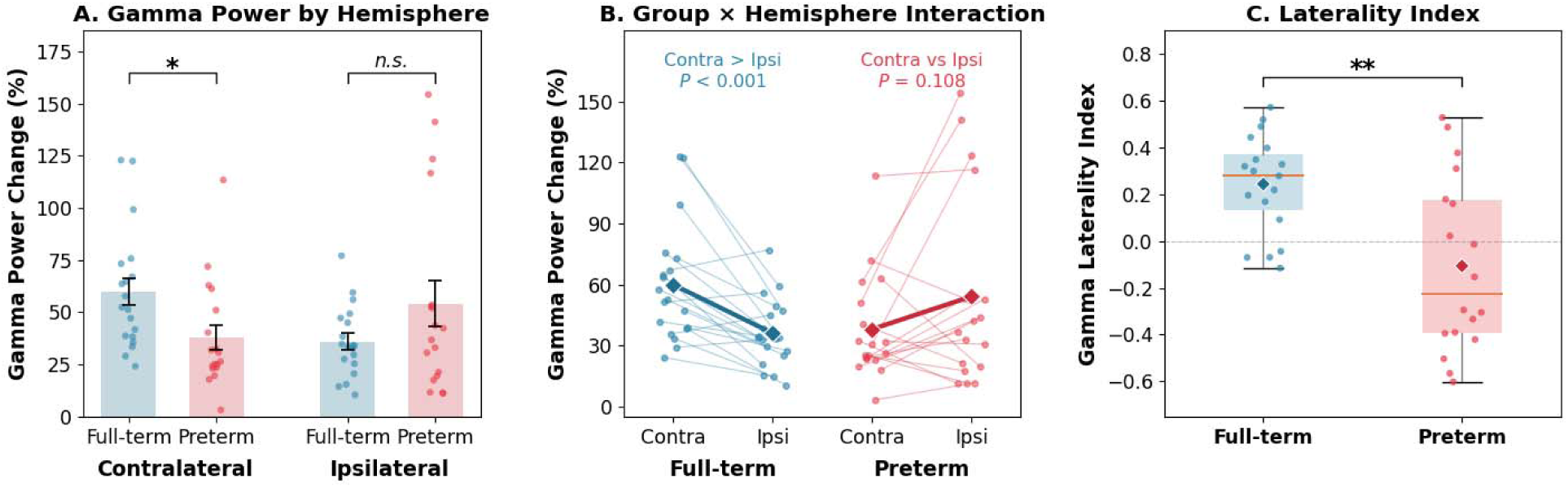
Group differences in M1 gamma power and hemispheric lateralisation. **(A)** Mean gamma power in contralateral and ipsilateral M1 for children born full-term and children born preterm. Contralateral gamma power was significantly higher in the full-term than preterm group (*t*(35) = 2.43, *P* = 0.020, Cohen’s *d* = 0.80), whereas the between-group difference in ipsilateral gamma power was not significant *(t*(21.1) = −1.55, *P* = 0.137, Cohen’s *d* = −0.52). **(B)** A two-way mixed analysis of variance revealed no significant main effect of group, *F*(1, 35) = 0.05, *P* = 0.823, partial _η_² = 0.001, and no significant main effect of hemisphere, *F*(1, 35) = 0.66, *P* = 0.421, partial _η_² = 0.019. There was a significant Group × Hemisphere interaction, *F*(1, 35) = 13.36, *P* = 0.000836, partial _η_² = 0.276. Follow-up paired-samples comparisons showed that gamma power was significantly greater in the contralateral than ipsilateral M1 in the full-term group, *t*(18) = 4.18, *P* = 0.0006, whereas no significant hemispheric difference was observed in the preterm group, *t*(17) = −1.70, *P* = 0.108. **(C)** Gamma laterality index, calculated as the normalised difference between contralateral and ipsilateral gamma power, was significantly greater in the full-term than preterm group (*t*(26.4) = 3.51, *P* = 0.002, Cohen’s *d* = 1.17). Points represent individual participants, and error bars in panel A indicate SD. Data are shown for 19 children born full-term and 18 children born preterm, with each data point representing one child. Two-tailed independent-samples *t*-tests were used for between-group comparisons, with Welch’s *t*-test used when the assumption of equal variances was violated. **Alt text:** Three-panel figure comparing motor-related gamma power and hemispheric lateralisation between children born full-term and preterm. Panel A shows higher contralateral M1 gamma power in the full-term group, while ipsilateral gamma power is more variable in the preterm group and does not differ significantly between groups. Panel B shows greater contralateral than ipsilateral gamma power in the full-term group but no significant hemispheric difference in the preterm group. Panel C shows a lower gamma laterality index in the preterm group, consistent with reduced contralateral dominance.

A two-way mixed analysis of variance revealed no significant main effect of group, *F*(1, 35) = 0.05, *P* = 0.823, partial _η_² = 0.001, and no significant main effect of hemisphere, *F*(1, 35) = 0.66, *P* = 0.421, partial _η_² = 0.019. There was a significant Group × Hemisphere interaction, *F*(1, 35) = 13.36, *P* < 0.001, partial _η_² = 0.276 (**Fig. 3B**). Follow-up paired-samples comparisons showed that gamma power was significantly greater in the contralateral than ipsilateral M1 in the full-term group, *t*(18) = 4.18, *P* < 0.001, whereas no significant hemispheric difference was observed in the preterm group, *t*(17) = −1.70, *P* = 0.108.

### Laterality of movement-related gamma oscillations

To further investigate hemispheric organisation of motor-related gamma activity, we calculated the laterality index as the normalised difference between contralateral and ipsilateral gamma power. The gamma laterality index was significantly higher in the full-term group than in the preterm group (0.24 ± 0.21 versus −0.11 ± 0.37; *t*(26.4) = 3.51, *P* = 0.002, Cohen’s *d* = 1.17; **Fig. 3C**). The positive mean laterality index in the full-term group was consistent with greater contralateral than ipsilateral gamma power on average, whereas the negative mean laterality index in the preterm group indicated relatively greater ipsilateral than contralateral gamma power on average.

Across the whole sample, greater contralateral gamma lateralisation was associated with higher K-ABC Hand Movements scores after FDR correction (ρ = 0.40, *P* = 0.014, *q* = 0.042, *n* = 37; **Fig. 4A**) and with earlier attainment of independent walking (ρ = −0.57, *P* < 0.001, *q* = 0.002, *n* = 35; **Fig. 4B**). No significant associations were observed with age at head control (ρ = −0.27, *P* = 0.107, *q* = 0.186, *n* = 37), age at sitting unsupported (ρ = −0.26, *P* = 0.124, *q* = 0.186, *n* = 36), VABS-II Gross Motor scores (ρ = 0.13, *P* = 0.440, *q* = 0.440, *n* = 36), or VABS-II Fine Motor scores (ρ = 0.15, *P* = 0.369, *q* = 0.440, *n* = 36).

**Figure 4.**
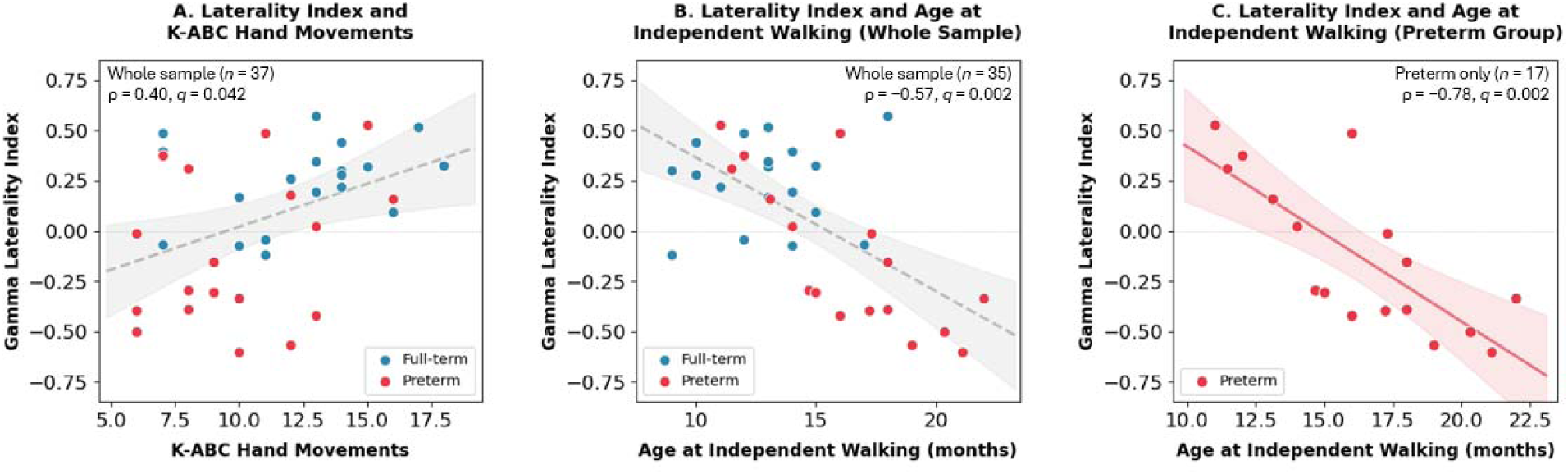
Associations between the M1 gamma laterality index and motor and developmental measures. **(A)** Spearman correlation between the M1 gamma laterality index and K-ABC Hand Movements scores across the whole sample (_ρ_ = 0.40, *P =* 0.014*, q =* 0.042, *n* = 37). **(B)** Spearman correlation between the M1 gamma laterality index and age at independent walking across the whole sample (_ρ_ = −0.57, *P* = 0.00035, *q* = 0.002, *n* = 35). **(C)** Spearman correlation between the gamma laterality index and age at independent walking in the preterm group (_ρ_ = −0.78, *P* = 0.00025, *q* = 0.002, *n* = 17). Lines indicate linear regression fits, and shaded areas indicate 95% confidence bands around the fitted regression lines for visualisation; statistical inference was based on Spearman’s rank correlations, with statistical significance determined using FDR-adjusted *q*-values. Each point represents one child. **Alt text:** Three scatterplots showing associations between the M1 gamma laterality index and motor-developmental measures. Panel A shows a positive association between gamma laterality and K-ABC Hand Movements scores across the whole sample. Panel B shows a negative association between gamma laterality and age at independent walking across the whole sample, indicating greater contralateral lateralisation with earlier walking. Panel C shows a negative association between gamma laterality and age at independent walking within the preterm group. Fitted regression lines and shaded confidence bands illustrate the direction of the associations.

Within the preterm group, greater contralateral gamma lateralisation was strongly associated with earlier attainment of independent walking after FDR correction (ρ = −0.78, *P* < 0.001, *q* = 0.002, *n* = 17; **Fig. 4C**). No significant associations were observed with age at head control (_ρ_ = −0.21, *P* = 0.410, *q* = 0.647, *n* = 18), age at sitting unsupported (ρ = −0.20, *P* = 0.431, *q* = 0.647, *n* = 18), K-ABC Hand Movements scores (ρ = 0.21, *P* = 0.396, *q* = 0.647, *n* = 18), VABS-II Gross Motor scores (ρ = 0.02, *P* = 0.944, *q* = 0.944, *n* = 18), or VABS-II Fine Motor scores (ρ = 0.13, *P* = 0.621, *q* = 0.745, *n* = 18).

## Discussion

In this study, we investigated the neurophysiological correlates of motor development following preterm birth. By combining MEG measures of motor-related gamma activity with perinatal and behavioural indices, we found that children born preterm showed longer response times and differences in cortical dynamics during voluntary movement. Reduced contralateral gamma power and reduced hemispheric lateralisation characterised the preterm group, while greater contralateral gamma lateralisation was associated with higher K-ABC Hand Movements score and earlier attainment of independent walking across the whole sample. These results suggest that preterm birth is associated with differences in cortical oscillatory activity and hemispheric organisation, and that motor-related gamma dynamics may reflect differences in motor-network organisation following preterm birth. A schematic summary of the main findings and their interpretation is provided in **Fig. 5**.

**Figure 5.**
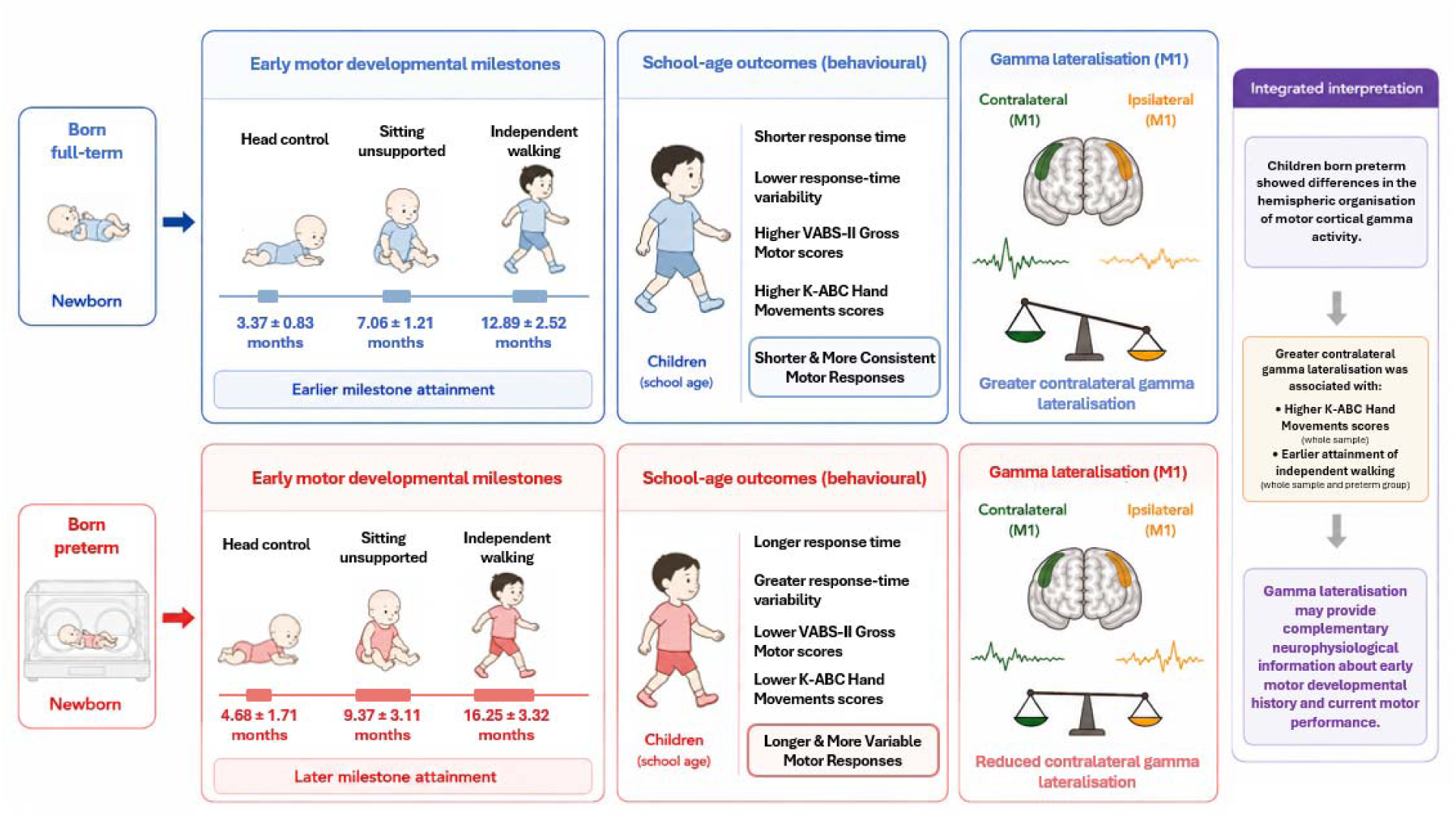
Schematic summary of motor development, behavioural performance, and movement-related gamma lateralisation in children born full-term and preterm. Children born preterm showed later attainment of early motor milestones, longer and more variable motor responses, and lower scores on selected motor measures at school age, together with reduced contralateral M1 gamma power and reduced contralateral gamma lateralisation. Greater contralateral gamma lateralisation was associated with higher K-ABC Hand Movements scores and earlier attainment of independent walking across the whole sample, with the association with independent walking also observed within the preterm group. Milestone ages are presented as mean ± SD and are expressed as chronological age for children born full-term and corrected age for children born preterm. The schematic summarises the observed findings and their integrated interpretation; the illustrated relationships should not be interpreted as evidence of causality. **Alt text:** Schematic comparing early motor development, school-age behavioural performance, and motor cortical gamma lateralisation in children born full-term and preterm. The upper row depicts children born full-term with earlier motor milestone attainment, shorter and more consistent responses, and greater contralateral gamma lateralisation. The lower row depicts children born preterm with later milestone attainment, longer and more variable responses, lower scores on selected motor measures, and reduced contralateral gamma power and gamma lateralisation. The integrated interpretation indicates that greater contralateral gamma lateralisation is associate with higher K-ABC Hand Movements scores and earlier independent walking, while presenting these relationships as associations rather than causal effects.

### Reduced Contralateral M1 Gamma Power Following Preterm Birth

A central finding was lower gamma power (70–90 Hz) in the contralateral M1 in children born preterm. Both groups showed movement-related gamma responses in this range, consistent with previous reports across children and adults.^19,21,23–25,44^ Motor cortical gamma oscillations have been linked to movement initiation and execution,^19,21,45,46^ while gamma-band synchronisation more broadly has been proposed to support cortical computation.^47^ Previous MEG studies have reported differences in cortical oscillatory activity and large-scale neural synchrony in preterm populations across cognitive^48,49^ and resting-state paradigms.^50^ These findings suggest that preterm birth may be associated with differences in cortical oscillatory coordination, consistent with the differences in movement-related gamma responses observed in the present study.

The reduced motor-related gamma power observed in children born preterm may reflect multiple interacting neurodevelopmental processes. First, gamma oscillations depend critically on the function and maturation of fast-spiking parvalbumin-positive GABAergic interneurons, which coordinate precise timing of excitatory activity within cortical microcircuits.^51,52^ Studies of preterm brain development have reported differences in GABAergic and interneuron maturation,^53,54^ providing a possible cellular basis for differences in gamma synchrony. Second, white matter and thalamocortical development may also be relevant. Imaging studies have shown differences in white matter microstructure and development^27,32^ and in thalamocortical connectivity^28,31^ following preterm birth. However, these mechanisms were not directly measured in the present study and therefore remain possible explanations rather than demonstrated mechanisms.

Importantly, the reduction in contralateral gamma power was observed at school age. This indicates that group differences in motor cortical activity are evident beyond infancy and early childhood, although the present cross-sectional data cannot determine whether reduced gamma activity represents a persistent developmental difference, variation in maturational timing, or a different developmental phenotype. Differences in gamma-band activity have been reported across a range of clinical conditions,^55^ including schizophrenia,^56,57^ autism spectrum disorder,^23,24^ attention-deficit/hyperactivity disorder,^58^ and Alzheimer’s disease,^59^ suggesting that variation in gamma activity is not specific to prematurity. Taken together, reduced motor-related gamma power in children born preterm may reflect differences in the maturation of cortical microcircuits and their interactions with developing white matter and thalamocortical networks. Paediatric MEG therefore provides a non-invasive approach for characterising neurophysiological differences in motor cortical function that may not be fully captured by behavioural measures alone.

### Lateralisation and Ipsilateral M1 Activity

The laterality index is a quantitative indicator of hemispheric dominance and reflects the relative balance between contralateral and ipsilateral activity.^43,60^ In the present study, children born preterm showed a marked reduction in hemispheric lateralisation of motor-related gamma oscillations. Compared with children born full-term, they exhibited lower contralateral gamma power and numerically greater ipsilateral gamma power, although the between-group difference in ipsilateral gamma power was not statistically significant, resulting in significantly lower laterality indices. The reduced laterality index therefore reflected the combination of reduced contralateral gamma power and relatively greater, although more variable, ipsilateral gamma power in the preterm group. Importantly, the laterality index was associated with selected developmental and motor measures across the whole sample, particularly K-ABC Hand Movements scores and age at independent walking. Greater contralateral gamma lateralisation was associated with higher K-ABC Hand Movements scores and earlier attainment of independent walking. The association with independent walking was also evident within the preterm group, suggesting that this relationship was not solely driven by the contrast between preterm and full-term groups. In contrast, the association with K-ABC Hand Movements scores was not significant within the preterm group and should therefore be interpreted more cautiously.

Movement-related gamma activity has been observed across childhood and adulthood, although how its hemispheric organisation changes across development remains incompletely understood.^19,21,23–25^ The association between gamma lateralisation and age at independent walking in the present study suggests that individual differences in the hemispheric organisation of motor-related activity may relate to variation in early motor development. The reduced lateralisation observed in the preterm group may therefore be consistent with differences in the developmental maturation of hemispheric motor organisation. This profile suggests that preterm birth may be associated with differences in the developmental organisation of neural mechanisms that support predominantly contralateral motor processing. The relatively greater ipsilateral activity may be consistent with reduced hemispheric specialisation, although the present data cannot establish whether this pattern reflects differences in maturational stage, compensatory recruitment, or other forms of motor-network reorganisation. More broadly, our findings complement recent functional MRI evidence showing persistent differences in task-evoked motor activation and large-scale functional network organisation in children born very preterm at a similar developmental stage.^33^ Together, these findings suggest that differences in the functional organisation of motor-related brain activity following preterm birth remain detectable at school age across complementary neuroimaging modalities. Differences in motor-network organisation have also been reported in other neurodevelopmental populations, including children with autism.^23,24,61^

Structural factors that influence interhemispheric communication may be relevant to this pattern. The corpus callosum is the largest white matter tract connecting neocortical regions across the hemispheres and enables interhemispheric information transfer and coordination.^62,63^ Within this structure, the callosal motor fibres specifically link the M1 of the two hemispheres, supporting bilateral motor integration.^64^ Previous studies have reported smaller callosal size^65–68^ and differences in callosal microstructure^27,29,30,69^ in individuals born preterm. These structural properties may have functional relevance, as callosal size and microstructural characteristics have been associated with motor performance and bimanual coordination.^64,70,71^ These findings raise the possibility that differences in callosal development may contribute to differences in interhemispheric communication and, in turn, to reduced motor cortical lateralisation and variation in motor performance following preterm birth. Although this interpretation is biologically plausible, callosal structure and connectivity were not directly assessed in the present study. The relationship between reduced gamma lateralisation and callosal development should therefore be tested directly using multimodal imaging in future work. More broadly, the associations between gamma lateralisation and motor-developmental measures should not be interpreted as evidence of a direct causal relationship. The whole-sample associations may partly reflect between-group differences, because the preterm group showed reduced lateralisation alongside later milestone attainment and lower scores on selected motor measures. The association with independent walking within the preterm group provides some evidence that this relationship is not entirely attributable to group status, although the relatively small sample limits the strength of this inference. Longitudinal studies are therefore needed to determine whether reduced gamma lateralisation precedes and relates to subsequent motor development or instead reflects adaptation within a different developmental trajectory.

### Linking Neural and Behavioural Findings: Longer Response Times in Children Born Preterm

Individuals born preterm frequently show differences in motor function.^17,72,73^ Even in the absence of cerebral palsy, these differences can include impaired fine and gross motor skills^72–74^ and difficulties in manual dexterity and visuomotor coordination.^17,74–76^ In the present study, children born preterm showed significantly longer button response times, with a mean difference of approximately 142 ms compared with children born full-term. This result indicates slower behavioural responses during the visuomotor task and may reflect differences in the timing of visuomotor processing and motor response initiation. Similar slowing has been widely reported in preterm populations across cognitive and sensorimotor domains, including reaction times in attention tasks,^5^ alertness responses,^77^ and magnitude comparison.^78,79^ A study of very preterm children aged 4–12 years also reported slower and more variable processing speed,^80^ and slower reaction times have been reported in young adults born very preterm.^81^

In the present study, mean response time was negatively associated, after FDR correction, with gestational age, birth weight, birth length, birth head circumference, and all three concurrent motor measures: K-ABC Hand Movements, VABS-II Gross Motor, and VABS-II Fine Motor scores. Thus, slower responses were associated with lower gestational age and smaller birth measures, as well as lower scores across multiple motor measures. For intra-individual response-time variability, associations with birth head circumference and K-ABC Hand Movements scores remained significant after FDR correction, whereas associations with birth weight and birth length did not. These secondary association analyses should be interpreted cautiously, as the intercorrelations among perinatal variables mean that their individual associations should not be interpreted as independent effects. Moreover, because these correlations were examined across the whole sample, they may partly reflect between-group differences and should not be interpreted as evidence of within-group relationships. Nevertheless, the overall pattern is consistent with previous evidence of differences in motor performance across childhood in individuals born very preterm or with very low birth weight.^11,17,18,82,83^

Together, these behavioural findings show that children born preterm had longer and more variable response times and lower scores on selected motor measures. The concurrent reductions in contralateral gamma power and hemispheric lateralisation provide complementary neurophysiological evidence of differences in motor-network organisation. Because the present analyses did not directly test associations between gamma and response-time measures, the findings do not establish a direct mechanistic link between differences in gamma activity and slower behavioural responses. The modest sample size and gestational-age profile of the preterm group may also limit the generalisability of the findings, particularly to children born late preterm.

## Conclusion

In conclusion, this study provides an MEG characterisation of motor cortical activity during voluntary movement in children born preterm. Children born preterm showed longer response times, reduced contralateral gamma power, and reduced hemispheric lateralisation, indicating differences in motor cortical dynamics during voluntary movement. Greater contralateral gamma lateralisation was associated with higher K-ABC Hand Movements scores and earlier attainment of independent walking across the whole sample, with the association with independent walking also evident within the preterm group. These findings suggest that differences in motor cortical function and hemispheric organisation following preterm birth are evident at school age. The observed reductions in gamma power and contralateral dominance may reflect differences in the maturation of motor cortical circuits and their interactions with developing white matter pathways and interhemispheric networks. In particular, reduced lateralisation may reflect differences in hemispheric specialisation of motor processing, potentially related to variation in callosal development. These mechanistic interpretations remain tentative because the underlying cellular and structural substrates were not directly measured. Together, these findings highlight the importance of considering both local cortical oscillatory activity and hemispheric organisation when investigating motor development following preterm birth. Movement-related gamma activity may provide complementary neurophysiological information about motor-network organisation alongside behavioural assessments. Future multimodal and longitudinal studies with larger samples will be important for clarifying the neural substrates associated with these oscillatory differences, determining how they evolve across development, and establishing their potential value for understanding individual developmental trajectories following preterm birth.

## Author contributions

J.Y. Data curation; Formal analysis; Methodology; Visualization; Writing—original draft; Writing—review & editing. S.S. Data curation; Formal analysis; Methodology; Visualization; Writing—original draft; Writing—review & editing. Y.M. Conceptualization; Investigation; Writing—review & editing. C.H. Investigation; Writing—review & editing. T.I. Investigation; Writing—review & editing. S.T. Investigation; Writing—review & editing. H.Y. Methodology; Writing—review & editing. K.T.J.C. Methodology; Writing—review & editing. A.W. Methodology; Writing—review & editing. K.Y. Investigation; Writing—review & editing. S.I. Investigation; Writing—review & editing. D.N.S. Investigation; Writing—review & editing. T.H. Investigation; Writing—review & editing. M.K. Conceptualization; Funding acquisition; Investigation; Project administration; Supervision; Writing—review & editing. Y.Y. Conceptualization; Data curation; Investigation; Methodology; Project administration; Supervision; Writing—review & editing. K.A. Conceptualization; Data curation; Formal analysis; Funding acquisition; Investigation; Methodology; Project administration; Supervision; Visualization; Writing—original draft; Writing—review & editing.

## Data availability

The data that support the findings of this study are available from the corresponding author upon reasonable request, subject to applicable ethical and institutional requirements. The data are not publicly available due to privacy and ethical considerations associated with paediatric neuroimaging and developmental data. Analysis code is available from the corresponding author upon reasonable request.

## Acknowledgements

The authors thank Sachiko Kitagawa and Yukiko Saotome for conducting the behavioural and MEG experiments, and Mutsumi Ozawa and Yoko Morita for preparing the experiments. The authors are especially grateful to the children and their parents who participated in this study. The authors acknowledge the use of ChatGPT (OpenAI) for language editing and refinement of manuscript wording. All AI-assisted content was reviewed and verified by the authors, who take full responsibility for the final manuscript.

